# LectinOracle – A Generalizable Deep Learning Model for Lectin-Glycan Binding Prediction

**DOI:** 10.1101/2021.08.30.458147

**Authors:** Jon Lundstrøm, Emma Korhonen, Frédérique Lisacek, Daniel Bojar

**Affiliations:** Department of Chemistry and Molecular Biology, University of Gothenburg, Gothenburg, Sweden. Wallenberg Centre for Molecular and Translational Medicine, University of Gothenburg, Gothenburg, Sweden; Swiss Institute of Bioinformatics, Geneva, Switzerland. Computer Science Department, UniGe, Geneva, Switzerland. Section of Biology, UniGe, Geneva, Switzerland

**Keywords:** glycobiology, machine learning, bioinformatics, carbohydrate, computational biology

## Abstract

Ranging from bacterial cell adhesion over viral cell entry to human innate immunity, glycan-binding proteins or lectins abound in nature. Widely used as staining and characterization reagents in cell biology, and crucial for understanding the interactions in biological systems, lectins are a focal point of study in glycobiology. Yet the sheer breadth and depth of specificity for diverse oligosaccharide motifs has made studying lectins a largely piecemeal approach, with few options to generalize. Here, we present LectinOracle, a model combining transformer-based representations for proteins and graph convolutional neural networks for glycans to predict their interaction. Using a curated dataset of 564,647 unique protein-glycan interactions, we show that LectinOracle predictions agree with literature-annotated specificities for a wide range of lectins. We further identify clusters of lectins with related binding specificity that are not clustered based on sequence similarity. Using a range of specialized glycan arrays, we show that LectinOracle predictions generalize to new glycans and lectins, with qualitative and quantitative agreement with experimental data. We further demonstrate that LectinOracle can analyze whole lectomes and their role in host-microbe interactions. We envision that the herein presented platform will advance both the study of lectins and their role in (glyco)biology.

## Introduction

Lectins, a class of glycan-binding proteins (GBPs), are present across all domains of life and play a fundamental role in a diverse range of biological functions by recognizing and binding specific carbohydrate structures on cell surfaces^[1]^. Examples of their importance in biology abound^[2]^. Following infection, activation of the complement pathway is regulated by mannose-binding lectin (MBL) recognizing mannose residues on the surface of pathogens. Homing of leukocytes during an adaptive immune response is coordinated through expression of selectins on the activated endothelium at the site of infection^[3]^. Viral and bacterial pathogens in turn use lectins to adhere to and infect target cells. The infectious activity of influenza viruses is mediated by hemagglutinin that binds sialic acid on the surface of cells in the upper respiratory tract. In fact, recognition of sialic acid in different contexts by different hemagglutinin sequences forms the basis for influenza host range^[4]^.

Lectins are often divided into many families based on sequence similarity and, consequently, common structural folds. Ligand binding is commonly enhanced by the presence of repeated glycan-binding domains (GBDs) and assembly into multimeric complexes, thus increasing the overall avidity for the cognate glycan^[5]^. While GBPs are found in all domains of life, the distribution of lectin families varies across taxonomic groups, suggesting the independent emergence of glycan-binding mechanisms during evolution^[6]^.

Glycans, the specific ligands of lectins, are composed of monosaccharides assembled into complex, branching structures and are present on the surface of all cells^[7]^. The composition and structure of glycans varies between cell types, species, and disease state, resulting in differential preferences for lectin interactions, particularly in complex samples^[8]^. Despite rapid methodological development in recent years, the experimental study of glycan structure and function remains limited compared to analogous investigations of DNA, RNA, and proteins, revolutionized by the emergence of affordable next-generation sequencing and high-sensitivity mass spectrometry. While sequential in nature, glycans are not included in the central dogma of biology and thereby do not benefit from sequencing-based approaches. Structural and functional investigations of glycans are further complicated by (i) non-templated synthesis regulated by subcellular localization and expression levels of glycosyltransferases, glycosidases, and nucleotide sugar substrates, and (ii) non-linear structures with multiple possible branch points and different linkages.

The intrinsic ability of lectins to selectively bind specific glycan motifs presents an opportunity to perform functional glycan investigation without the need for explicitly determining monosaccharide sequences. In experimental settings, lectins are used e.g., for characterizing molecular interactions, investigating cell identities, and mitogenic stimulation^[1]^. In a clinical context, the glycan-binding properties of lectins can be exploited in the development of therapeutic strategies, e.g., neutralization of bacterial toxins, or interfering with cellular adhesion, thus preventing viral infection. However, only a fraction of the known lectins has been fully characterized regarding glycan-binding specificities, limiting the readily available resources for experimental studies of glycan function.

Recently, ~ 1,000,000 putative lectin sequences were identified from the UniProt datasets from > 24,000 species, providing a database of possible lectins, e.g., for use in future biochemical analysis and therapeutics development^[9]^. In the past two decades, glycan-binding profiles of lectins have been mapped using glycan arrays. Here, hundreds of distinct glycans structures are immobilized on a glass surface in order to quantify the glycan-binding ability of a specific protein sample. While providing accurate binding specificities, the individual experimental investigation of thousands of lectins using glycan arrays is currently not feasible. However, computational predictions informed by machine learning models could narrow down the experimental search space.

Machine learning algorithms provide predictions for unseen input data, based on relationships learned from labelled training data. Deep learning structures such algorithms in layers, enabling the identification of salient features without having to explicitly designate them. This creates a model that can be optimized and ultimately outperformed human ability in tasks such as language processing or computer vision^[10,11]^. Recent advances in deep learning have provided neural net architectures capable of solving highly complex biological problems. Accurately predicting protein structure from amino acid sequence, seeming like an insurmountable task a few years ago, is now readily available with tools such as AlphaFold2 and RoseTTAFold^[12,13]^. In glycobiology, deep learning has recently enabled new analyses of sequence-function relationships^[14]^. Based on this, we developed SweetNet^[8]^, a graph convolutional neural network method that learns glycan representations by taking their branching structures into account.

Given the obvious benefits of a model predicting protein-glycan interactions, this challenging task would be a valuable application of models such as SweetNet. Previous efforts in predicting lectin-glycan interaction perform reasonably well in recapitulating already-determined experimental data^[15,16]^ but lack scalability and the possibility of providing predictions for novel lectins, which is essential for making more glycan-binding proteins available and thus empowering future investigations. Further, previous approaches have, at least in part, lacked model interpretation – learning what the model learned – as well as model application to understanding the manifold roles of lectins in biology.

Combining branch-aware glycan representations of a SweetNet-type model with a transformer-based protein sequence language model, we propose a generalizable computational framework for lectin-glycan interaction predictions. Our model, LectinOracle, accurately predicts Z-score transformed relative fluorescence units for lectin-glycan interactions, showing significant correlation with data from various custom glycan arrays not included during training and agreement with literature on newly characterized lectins. Based on predicted glycan-binding specificity of characterized and uncharacterized lectins, we suggest a lectin classification system – aided by sugar specificity in addition to protein folds – that spans taxonomic groups, simplifying the task of selecting suitable lectins for experimental purposes. By providing open access to LectinOracle, we aim to provide necessary tools for expanding our understanding of lectin-glycan interactions in diverse topics such as disease susceptibility, host-microbe interactions, agriculture, and (auto)immune disease.

## Results

### Developing a deep learning-based model to predict lectin-glycan interactions

In order to develop a model to predict and analyze protein-glycan interactions, we reasoned a setup was necessary that considered both protein and glycan information to arrive at a binding prediction. This was based on our vision to use the resulting model to extrapolate to interactions of new lectins as well as new glycans. We used protein sequences as inputs for our model rather than protein structures, as the amount of data available for the former far surpasses the latter. Importantly, sequence-based data contain evidence for glycans and proteins that do not interact, a crucial type of data that is lacking from protein-glycan co-crystallizations which only depict successful interactions. Further, protein-glycan co-crystallizations typically contain only very short glycan fragments, represent non-quantitative data, and, due to the molar excess of glycan fragments during crystallization and the absence of competing glycans, might not constitute physiologically relevant interactions. Finally, working with sequences also ensures that our model can readily be extended to uncharacterized lectins, as sequences are substantially easier to acquire than structures. To train such a model, we constructed a comprehensive dataset of 564,647 unique protein-glycan interactions from 3,328 glycan microarray experiments (Table S1-2), from the Consortium for Functional Glycomics as well as the Carbohydrate Microarray Facility of Imperial College London. Our dataset consisted of 1,392 lectins, including plant, fungal, bacterial, viral, and animal lectins (Figure 1B), as well as 927 glycans (Figure 1C). On these data, we trained deep learning-based models, predicting Z-score transformed relative fluorescence units (representing binding) based on protein and glycan sequences.

**Figure 1.**
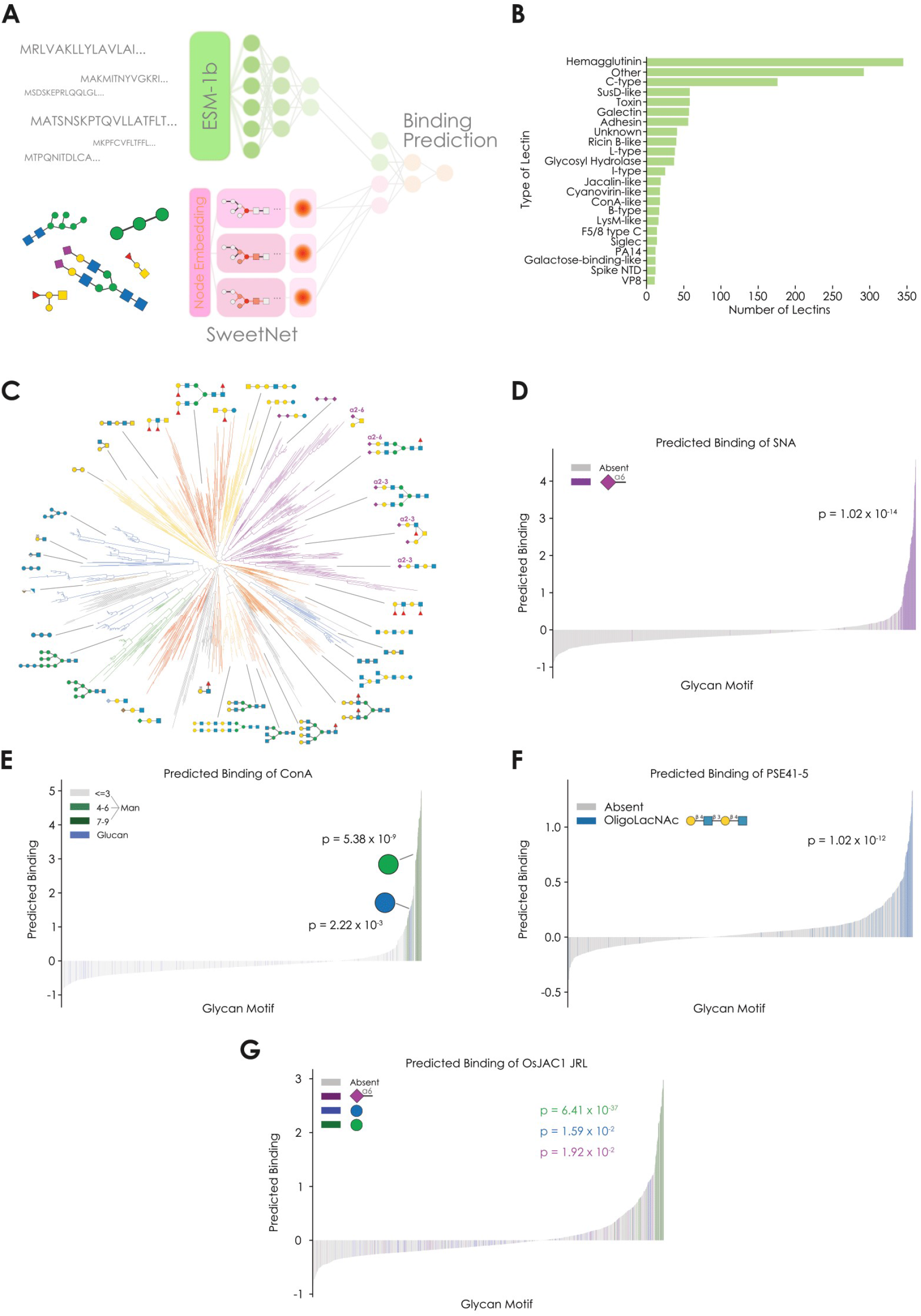
Predicting lectin binding specificity with deep learning. **A)** Scheme of the deep learning model LectinOracle, which analyzes protein sequences via a pre-trained transformer-based model (ESM-1b) that is further fine-tuned, and glycan sequences via a graph convolutional neural network (SweetNet). Results from both arms of the model are concatenated and used to predict lectin-glycan binding. **B)** Dataset Proportions. Shown is the composition of our lectin-glycan dataset for training LectinOracle, with the proportions of each lectin class depicted. **C)** Glycan clusters in dataset. The diversity and relatedness of glycans in our dataset is shown via a dendrogram obtained by neighbor joining of the representations learned by the SweetNet component of LectinOracle. The dendrogram is visualized via the Interactive Tree of Life v5.5 software^[19]^ and annotated with glycan groups. **D-E)** Analysis of known lectins via LectinOracle, with the example of SNA (**D**) and ConA (**E**). For a range of glycan motifs, a trained LectinOracle model was used to predict their binding to the lectins SNA (D) or ConA (E), with their literature-annotated binding motifs colored. **F-G)** Analysis of uncharacterized lectins, with the example of PSE41-5 (**F**) and the jacalin-related domain from OsJAC1 (**G**). Motifs which were identified to be enriched in predicted bound glycans are colored. Motif enrichment was tested via one-sided Wilcoxon signed-rank tests, with the p-value shown in each panel.

To analyze glycan sequences, we used the state-of-the-art SweetNet architecture, as it has been shown to generally outperform alternative methods and to efficiently scale with glycan diversity^[8]^. SweetNet comprises a graph convolutional neural network that was designed to accommodate the branching structures of glycans. In general, a deep learning-based glycan analysis module learns similarities between glycan motifs and can more easily generalize to new motifs than a discretized, motif counting-based approach that is more suited to data analysis than prediction^[14]^.

For analyzing protein sequences, we evaluated different approaches. Previous work used recurrent neural networks (RNNs), a type of language model, to predict the interaction of viral hemagglutinin proteins with glycans^[8]^, analyzing receptors for influenza virus. Yet the analysis of protein sequences via deep learning has greatly progressed and transformer-based models have been shown to outpace RNNs for purposes such as predicting protein function^[17,18]^. Therefore, we chose protein representations learned by ESM-1b (Evolutionary Scale Modeling 1b)^[18]^, a 650 million parameter model trained on the entirety of UniRef50, as the input for our model, so that this rich representation could be further fine-tuned for the task presented here.

After processing both protein and glycan sequences, our model concatenated representations learned for both interaction partners and used this information to predict protein-glycan binding via a fully connected neural network module. We evaluated different variations of this model scheme (Table 1) and concluded that the variant with a fine-tuned ESM-1b module for protein sequences, a SweetNet-based module for glycan sequences, and a subsequent fully connected neural network with multi-sample dropout and sigmoid output scaling resulted in the best empirical performance. We therefore based all further analyses on this model, which we have named LectinOracle (Figure 1A), that has been trained on a wide range of different lectin classes (Figure 1B).

**Table 1:** Selecting an architecture for a model predicting protein-glycan interactions. For the task of predicting protein-glycan interactions, we trained deep learning models with varying architectures to identify a suitable model for this study. In this table, we note the differences in the various models in three modules, the arm analyzing protein sequences, the arm analyzing glycan sequences, and the downstream module combining protein and glycan information for binding prediction. Mean values from five independent training runs (Table S3) are provided for the mean squared error (MSE) and mean absolute error (MAE) on a separate test set. For each metric, the superior value is bolded.

| **Protein Arm** | **Glycan Arm** | **Combined Head** | **MSE** | **MAE** |
| --- | --- | --- | --- | --- |
| RNN | SweetNet | Fully connected +<br>Multi-sample dropout +<br>Sigmoid | 0.9925 | 0.501 |
| ESM-1b | SweetNet | Fully connected +<br>Multi-sample<br>dropout +<br>Sigmoid | 0.7475 | 0.4276 |
| ESM-1b + fine-tune | SweetNet | Fully connected +<br>Sigmoid | 0.7375 | 0.4238 |
| ESM-1b + fine-tune | SweetNet | Fully connected +<br>Multi-sample<br>dropout | 0.7415 | 0.4301 |
| ESM-1b + fine-tune | SweetNet | Fully connected +<br>Multi-sample<br>dropout +<br>Sigmoid | **0.7283** | **0.4137** |

Next to predicting protein-glycan interactions, a trained LectinOracle model can also be used to retrieve representations – learned similarities – for both proteins and glycans. These representations can be used to cluster sequences, such as the glycan sequences in our dataset (Figure 1C). Importantly, “similarities” are task-dependent and therefore should, in this case, reflect similarities in binding behavior. While tasks such as predicting the taxonomic origin of a glycan typically result in glycan representations clustered by class (N-linked, O-linked, etc.)^[8,14]^, here we observe a clustering that spans classes and is largely influenced by terminal glycan motifs (e.g., α2-3 linked Neu5Ac, α2-6 linked Neu5Ac, terminal GalNAc). Since these terminal motifs are crucial for determining lectin-binding^[20]^, we concluded that LectinOracle seemed to have learned to extract relevant information from glycans to predict protein-glycan interactions.

We then evaluated the performance of a trained LectinOracle model by analyzing predictions for well-characterized lectins. First, we chose the lectin SNA (*Sambucus nigra* agglutinin), as it has a well-defined binding specificity for α2-6 linked Neu5Ac^[21]^. For a range of 1,578 glycan motifs occurring in our dataset (see Methods for details), we retrieved binding predictions for SNA from LectinOracle (Figure 1D). Among the predictions, we observed a striking enrichment for α2-6 linked Neu5Ac-containing motifs, which was highly significant based on a Wilcoxon signed-rank test (p = 5.38 x 10^-9^). In contrast to the highly specific binding specificity of SNA, we investigated the binding behavior of the lectin AAL (*Aleuria aurantia* lectin) and confirmed the broad recognition of fucose residues^[22]^ in various linkages (Figure S1A). We additionally analyzed the lectins SBA (soybean agglutinin), from *Glycine max*, and HPA (*Helix pomatia* agglutinin) and demonstrated that LectinOracle correctly learned their preference for terminal GalNAc residues^[23,24]^ (Figure S1B-C).

Next, we analyzed Concanavalin A (ConA), from the jack-bean *Canavalia ensiformis*, which is known to bind to mannose-rich glycans^[25]^. We observed an overwhelming enrichment of high-mannose structures in motifs that were predicted to be bound by ConA (Figure 1E), again confirming its literature-annotated binding specificity. Additionally, we identified a weaker binding preference for glucan motifs, which is consistent with reports that ConA can bind glucose-rich sequences^[26]^, such as from fungi. This clear separation between dominant (mannose-rich) and secondary (glucose-rich) binding motif that we observed for ConA suggested that the absolute predicted binding by LectinOracle scales, to a certain extent, with affinity or binding strength, which we also explored further in later sections. We indeed observed that, for both SNA and ConA, the binding prediction, on average, increased with a higher number of binding motifs in a glycan (Figure S2).

We also leveraged the generalizability of LectinOracle to investigate predicted binding motifs for lectins that are only coarsely characterized, or even entirely uncharacterized. One example for this is the lectin PSE41-5, identified by reverse vaccinology from *Pseudomonas aeruginosa*^[27]^, which has been shown to bind to terminal beta-linked galactose. With LectinOracle, we confirmed that type II LacNAc structures, with a terminal beta-linked galactose, were strongly enriched among the predicted binding motifs (Figure 1F). Applied to the relatively uncharacterized jacalin-related domain from OsJAC1, a lectin from the important food crop *Oryza sativa* (Asian rice), LectinOracle predicted binding predominantly to mannose-containing motifs yet also secondary binding to glucose and, specifically, Neu5Ac(α2-6)-containing motifs (Figure 1G). Recent studies with mono- and disaccharides have indeed shown binding to mannose and glucose for this domain^[28]^, whereas sialic acid was not tested for binding. Interestingly, the original jacalin, derived from jackfruit, has been shown to be capable of binding to Neu5Ac^[29]^, lending further support to our analyses.

We also predicted the binding specificity of OTV1_139, a hypothetical protein from *Ostreococcus tauri* virus 1^[30]^ that contains a CBM47 (carbohydrate-binding module 47) domain, which has been shown to bind fucose^[31]^. LectinOracle, applied to OTV1_139, also revealed a preference for binding fucose (Figure S1D). We next investigated two more lectins, *Arundo donax* lectin (ADL) and hypothetical cytosolic protein 031524 from *Bacillus subtilis* (YesU). For ADL, LectinOracle correctly inferred the activity of its chitin-binding domain^[32]^ and predicted motifs such as chitotriose as the top binders (Figure S1E). YesU has been characterized to prefer fucosylated glycans^[33]^ and LectinOracle also predicted fucosylated glycan motifs to be strongly enriched for binding (Figure S1F), including fucosylated motifs such as Lewis X. These case studies involving lectins outside our dataset emphasize that LectinOracle can be used generally to further probe the glycan-binding specificity of lectins, both already characterized as well as uncharacterized.

### Analyzing lectins with LectinOracle reveals lectin clusters with shared binding patterns

The categorization of lectins into different classes, such as galectins^[34]^ or LysM-like lectins^[35]^, has aided the understanding of lectin interactions on a systematic level. Often, lectins are categorized based on shared structural characteristics, such as a common fold^[9,36]^. However, structural similarity does not always lead to similar overall binding preferences and distinct protein folds, with shared binding specificity, have evolved independently^[6]^. We hypothesized that a lectin classification that also factored in (predicted) binding specificity would improve on this structural approach and lead to classes that would be immediately useful to researchers who use lectins in their work and rely on their binding specificity rather than their structure.

When coloring lectins based on the disaccharide that is the top prediction from LectinOracle, we overall did not observe meaningful binding specificity clustering in the ESM-1b representation, that solely relies on sequence similarity (Figure 2A). Possible exceptions to this were Neu5Ac-binding lectins that exhibited high sequence similarity and were prevalent in our dataset, such as the influenza virus hemagglutinins^[37]^. However, once we plotted the lectin representations learned by LectinOracle (sequence similarity + binding specificity), we immediately noticed several distinct lectin clusters that seemed to share binding patterns (Figure 2B).

**Figure 2.**
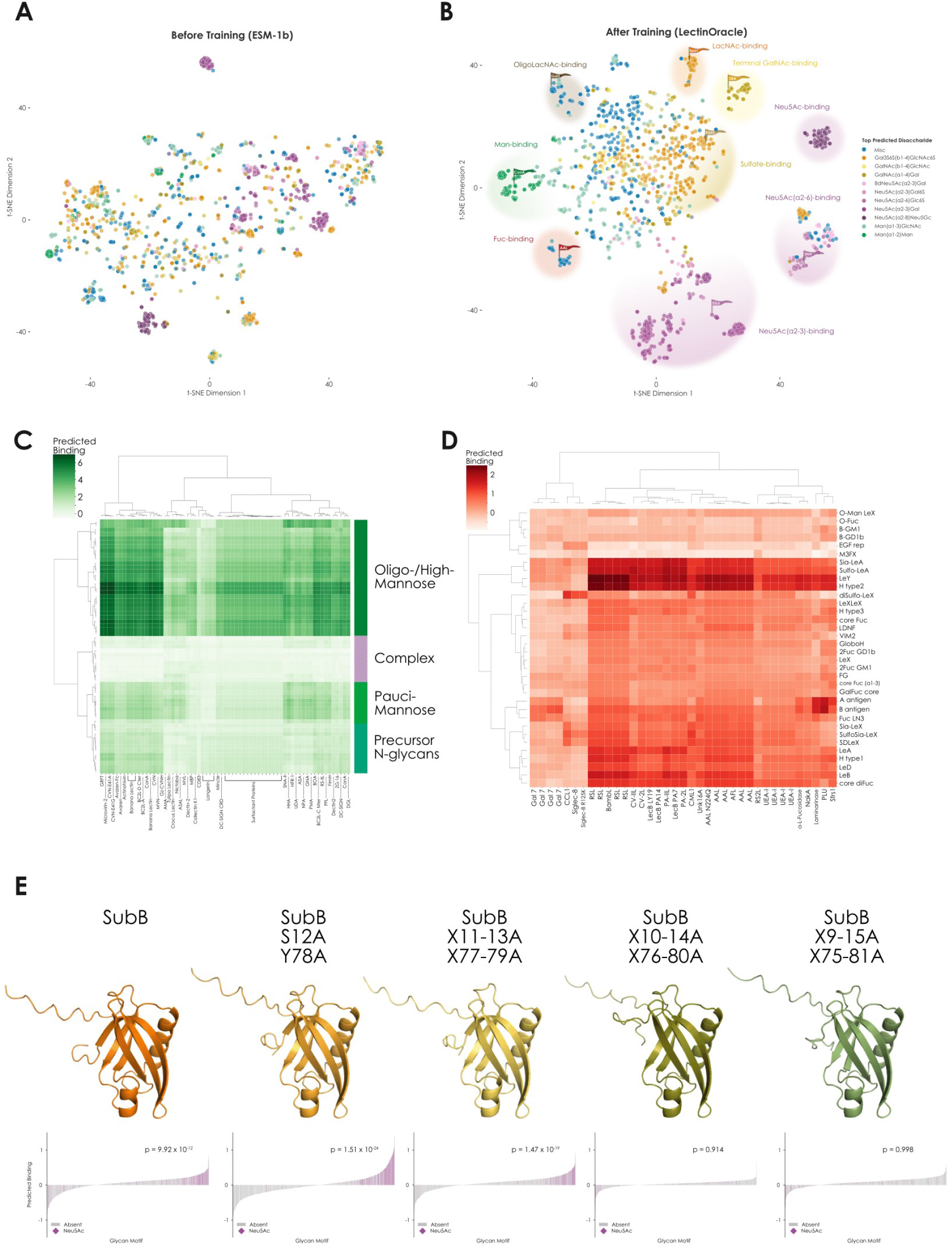
Clustering lectins based on learned binding motifs. **A-B)** Lectins clustered based on sequence similarity or binding specificity. Learned representations from the pre-trained ESM-1b model (**A**) or a trained LectinOracle model (**B**) were extracted for each lectin and are shown via t-SNE^[38]^, colored by the top predicted disaccharide binding motif. Cluster binding specificities are annotated and representative lectins are labeled. **C)** Characterization of mannose-binding lectins. We selected the group of lectins with ‘Man(a1-2)Man’ as the top-predicted disaccharide and used LectinOracle to predict their binding to a range of mannose-containing glycans. Then, we performed hierarchical clustering using Ward’s method^[39]^. **D)** Characterization of fucose-binding lectins. After selecting lectins with fucose within their top predicted disaccharide, we predicted their binding to a range of fucose-containing motifs from the academic literature and filtered out lectins which did not have at least a predicted binding of one to any of these motifs, depicting the rest as a hierarchical clustering based on Ward’s method. **E)** LectinOracle predictions are sensitive to amino acid substitutions in the lectin binding pocket. For SubB and various alanine-substitution mutants, we predicted their glycan-binding behavior with a range of glycan motifs. One-sided Wilcoxon signed-rank tests were used to ascertain significant enrichment for predicted Neu5Ac binding in each case. Protein structure predictions made with AlphaFold2 are shown for the wild-type protein and each mutant.

Most strikingly, we observed a mannose-binding lectin cluster that was not apparent in the clustering based on sequence similarity. This cluster contained well-known mannose-binding lectins, such as plant-derived ConA and the Banana lectin from *Musa acuminata* as well as mammalian lectins, such as the human MBL2 (Mannose Binding Lectin 2). While most clusters exhibited a dominant disaccharide that characterized cluster binding, fucose-binding lectins demonstrated a pronounced diversity in their exact fucose specificity between lectins, with no single overarching disaccharide despite forming a cluster in our representation. With regards to sialic acid-binding, we could annotate multiple clusters, with specificity for α2-3 linked Neu5Ac, α2-6 linked Neu5Ac, and Neu5Ac without linkage-preference, respectively. These clusters contained the expected avian (α2-3 linked Neu5Ac) as well as mammalian influenza virus hemagglutinins (α2-6 linked Neu5Ac)^[40]^. Yet we also identified other lectins in these clusters that were less obvious via sequence similarity, such as *Staphylococcus aureus* superantigen-like 6 (SSL6) or *Escherichia coli* heat-labile enterotoxin for the α2-3 linked Neu5Ac cluster and SNA or *Polyporus squamosus* lectin (PSL) in the α2-6 linked Neu5Ac cluster. This clustering beyond mere sequence similarity, which was enabled by LectinOracle, holds promise for re-classifying existing lectins as well as annotating and discovering new lectins that could be used as research probes or shed light on biological phenomena.

Next, we sought to further characterize the clusters unveiled by the similarities learned by LectinOracle (Figure 2B) and thereby propose a more useful characterization of lectins, not only informed by sequence similarity but additionally by binding specificity. We first analyzed the cluster containing mannose-binding lectins, which not only cleanly separated different classes of mannose-containing glycans based on their predicted binding but also revealed lectin sub-groups in this cluster that exhibited slightly different binding preferences (Figure 2C). One cluster, containing cyanovirin-like lectins, such as microvirin or cyanovirin, as well as the jacalin-like Banana lectin, was predicted to bind especially well to oligo- and high-mannose glycans. Similar to a dose-response relationship, lectins with a higher predicted binding to oligo-/high-mannose structures also exhibited, albeit lower, residual predicted binding to pauci-mannose structures (3-4 mannoses). This high-level characterization might serve as a guideline for researchers choosing a lectin that is best suited to their experimental needs.

Analogously, we analyzed fucose-binding lectins, based on widely known fucose-containing motifs from the academic literature (Figure 2D). Our first observation was that, while galectin 7 and siglec 8 were filtered together with the fucose-binding lectins, the motif heatmap revealed that they were only predicted to substantially bind to sialylated / sulfated motifs (siglec 8) or motifs containing LacNAc structures (galectin 7), not to other fucosylated structures. The remaining block of lectins represented the “true” fucose-specific lectins. Here, we also observed a dose response, in that difucosylated motifs (Lewis B, Lewis Y, difucosylated N-glycan cores, etc.) were, on average, predicted to be bound stronger than monofucosylated motifs. With analyses such as these, our platform constitutes a rapid means to characterize and compare a large set of lectins with regards to their binding specificity to identify sub-clusters as well as the most appropriate lectin reagents for detecting glycan motifs.

We were also intrigued to observe two distinct clusters for lectins predicted to bind the N-Acetyllactosamine (LacNAc) motif that is near-ubiquitous in animals (Figure 2B; light brown and dark brown). We therefore decided to identify what distinguished lectins from both clusters. As type I and type II LacNAc can occur in multiple forms (LacNAc, OligoLacNAc, PolyLacNAc), we used our trained LectinOracle model to predict the binding of all lectins in our dataset to one, two, or three repeats of type II LacNAc (Gal(β1-4)GlcNAc(β1-3); Figure S3). When coloring all lectins according to their predicted binding, we observed that one of the two clusters started out with strong predicted binding to 1xLacNAc yet seemed to lose predicted binding with 3xLacNAc. This cluster was enriched in epithelial adhesins from the fungus *Candida glabrata*^[41]^, The other cluster, however, increased in predicted binding with increasing number of LacNAc repeats. Here, we found an enrichment for galectins from various species. This led us to the conclusion that the first cluster constituted LacNAc-binders while the second cluster was rather characterized by oligoLacNAc-binding. This also implied that LectinOracle was sensitive to the number of repeats for a glycan motif, which we already indicated with the example of SNA and ConA above (Figure S2).

Beyond short di- or trisaccharide motifs, many larger motifs, such as the Lewis structures^[42]^, are widely known and relevant in various biological contexts. We therefore engaged in an analysis to find out whether we could identify clusters of lectins binding to these motifs (Figure S4). While we could pinpoint distinct regions for many motifs in the lectin space learned by LectinOracle, clustering in most cases could be explained by smaller motifs within larger motifs such as the Lewis series. The presence of fucose in Lewis structures and blood group epitopes led to activity in the fucose-binding cluster, motifs such as SialylLewis X added predicted binding from Neu5Ac-binding lectins, and the blood group A antigen included activity from the terminal GalNAc-binding cluster. Antibodies with high specificity for a defined large motif notwithstanding, most lectins in our dataset seem to be well-characterized on the disaccharide level.

Having established the sensitivity of LectinOracle to minute, monosaccharide-level changes in glycans, we then set out to analyze model sensitivity towards the lectin sequence. In other words, did LectinOracle learn a broader domain - motif relationship or a more fine-grained sequence / binding pocket - motif association? For this, we turned to *Escherichia coli* Subtilase cytotoxin subunit B (SubB), a protein outside our dataset and part of a bacterial AB5 toxin that binds sialic acids with two key amino acid residues, S12 and Y78^[43]^. We analyzed the glycan binding of SubB with LectinOracle and indeed observed significant enrichment of Neu5Gc- and Neu5Ac-containing moieties among the predicted binding glycans (Figure 2E, Figure S5). We note that, while we did observe a statistically significant binding prediction for Neu5Gc binding (Figure S5), due to a relative scarcity of Neu5Gc-containing glycan motifs on the arrays used for training LectinOracle we were unable to quantify any differential preference of Neu5Gc over Neu5Ac. Interestingly, we found that the binding predictions for SubB are highly sensitive towards amino acid substitutions in the glycan-interacting region of the protein, with Neu5Ac/Gc enrichment being completely abolished upon substituting five residues around each key amino acid with alanine (Figure 2E, Figure S5), while the substitutions did not substantially impact the protein structure as predicted by AlphaFold2^[12]^. Overall, these results demonstrate that LectinOracle, without any prior knowledge or being trained for this, is sensitive towards mutations of amino acids in the ligand binding pocket of SubB.

### LectinOracle predictions match independent experimental observations qualitatively and quantitatively

The gold standard for validating machine learning models is to compare their predictions to independent experimental observations. Not only does this procedure allow for the evaluation of the quality of a model but it also showcases the range of scenarios where a model can be applied with sufficient accuracy, as model predictions typically deteriorate the further a scenario is removed from the training data.

As a first validation, we compared our predictions to co-crystallized lectin-glycan structures found in UniLectin3D^[44]^. For this, we used the lectin and the glycan in the complex as input to LectinOracle to see whether this binding would have been predicted. For lectins that were part of our training set, LectinOracle achieved good predictive accuracy for lectins from all taxonomic groups (Figure S6A) and nearly all lectin classes (Figure S6B). When including all lectins, we did observe lower performance for archaeal lectins (absent from our dataset) and viral lectins (Figure S6C-D). As mentioned before, while offering valuable mechanistic insights into protein-glycan interactions, co-crystallized lectin-glycan structures should not be seen as the gold standard of predicting lectin-glycan binding due to potential limitations. We were nonetheless satisfied that LectinOracle, on average, predicted the interactions of the majority of lectins on UniLectin3D.

To circumvent potential issues of crystal structures, we also used experimental data from a range of customized glycan arrays that were not part of the CFG or Imperial College database to validate LectinOracle. These arrays contained glycans as well as lectins that were new to LectinOracle and offered a rich source for validating our predictions. First, we used the recently published oligomannose array for this purpose^[45]^. Here, investigators assayed a variety of subtly different oligomannose glycans for their binding to various lectins. The slight changes in glycan structure, which nonetheless led to appreciable differences in their binding behavior, made this a particularly challenging case study for our model. When thresholding our predictions and the observed relative fluorescence units (i.e., separating glycans into “bound” / “not bound” for a given lectin), we achieved predictive accuracies of ~70 - 90% for plant and bacterial lectins using LectinOracle (Figure 3A, Figure S7A). With the exception of Dectin-2, we observed similar results for mammalian lectins on this array (Figure S7B-D). We note that for the antibodies tested on this array, LectinOracle did not yield correct predictions (Figure S7C-D). This is likely due to two factors: (i) the antibodies are highly specific, with a low single-digit number of bound glycans on the oligomannose array that are easier to miss entirely and (ii) our training set lacks data for antibodies, so LectinOracle was never trained to be able to predict antibody-glycan binding and should not be used for this purpose.

**Figure 3.**
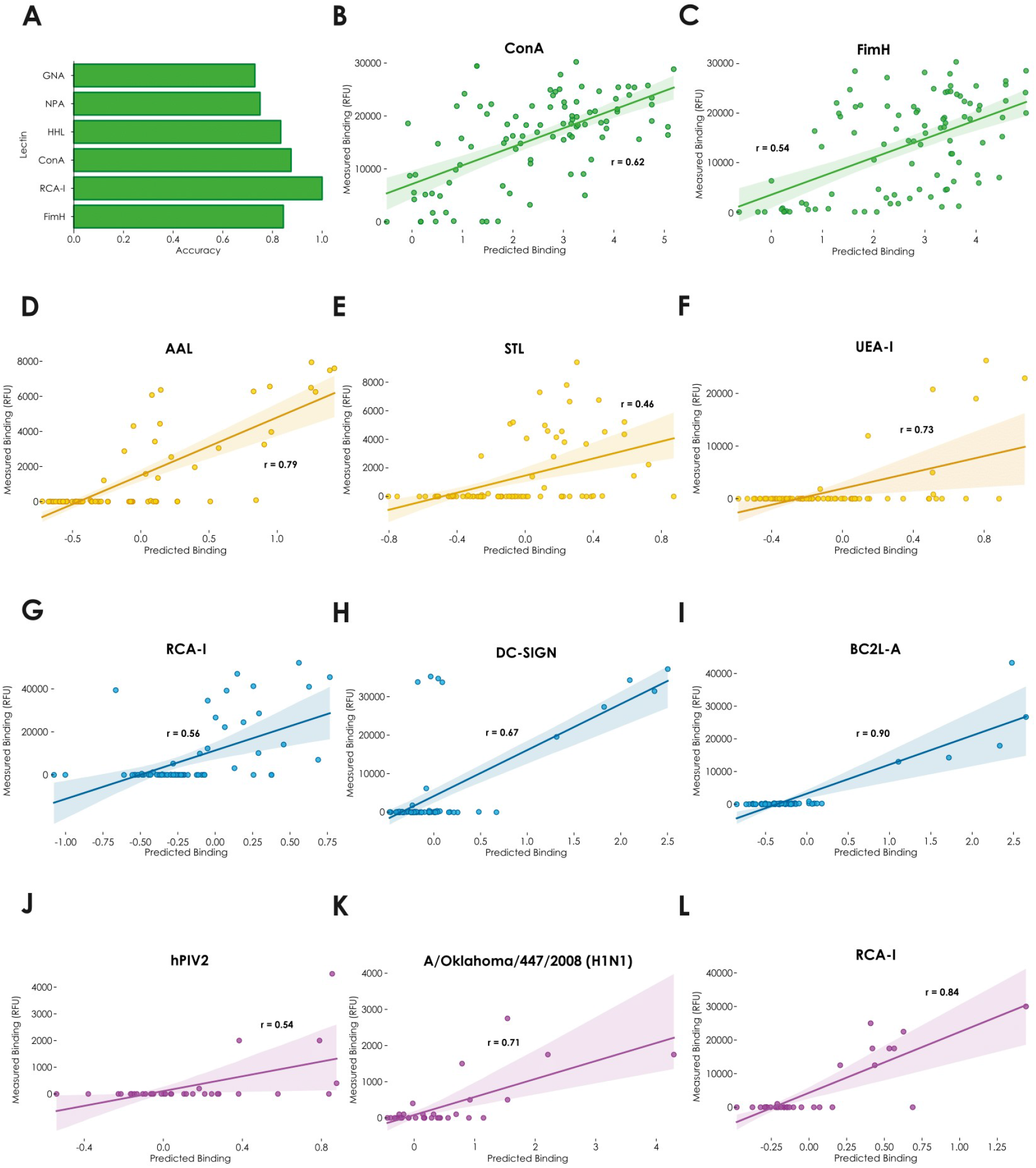
LectinOracle predicts binding of lectins to a wide range of different glycan arrays. **A)** Accuracy of LectinOracle predictions for lectins tested on the oligomannose array. For each lectin-glycan pair, we assigned it the label “bound” or “predicted bound” if the observed relative fluorescence units (RFU) were at least 10% of the maximum RFU or if the predicted binding was at least 0.5, respectively. The agreement of experiment and prediction is shown in terms of accuracy (precision can be found in Figure S7A). **B-C)** Correlating experimentally observed binding with predictions for the lectins ConA (B) and FimH (C) tested on the oligomannose array. **D-F)** Correlating experimentally observed binding with predictions for the lectins AAL (D), STL (E), and UEA-I (F) on the mucin *O*-glycan array. **G-I)** Correlating experimentally observed binding with predictions for the lectins RCA-I (G), DC-SIGN (H), and BC2L-A (I) on the microbe-focused array. **J-L)** Correlating experimentally observed binding with predictions for the lectins hPIV2 (J), A/Oklahoma/447/2008 (H1N1) (K), and RCA-I (L) on the sialic acid array. All correlations between experimental data and predictions were done via fitting a linear regression and r represents Pearson’s correlation coefficient.

Encouraged by these results, we hypothesized that the relative predicted binding from LectinOracle could be informative beyond classifying glycans into “bound” / “not bound”, as LectinOracle was trained on quantitative binding data. Therefore, for each glycan, we correlated predicted binding and experimentally measured binding for a range of lectins (Figure 3B-C, Figure S7B), to see whether we could predict quantitative binding, a substantially more challenging task. For lectins on the oligomannose array, we indeed achieved moderate correlations (defined as a Pearson’s correlation coefficient between 0.5 and 0.7) between our predictions and the experimental results, demonstrating that our predictions contain quantitative information about protein-glycan interactions, even in a different array setting than LectinOracle was trained on.

To show that this constitutes a general property of LectinOracle and to test the generalizability of its predictions in different contexts, we then went on to validate our predictions on other glycan arrays. First, we tested the recently reported mucin *O*-glycan array^[46]^, in which 83 *O*-linked glycans with various modifications were tested against several lectins. In this context, with very different glycans compared to the oligomannose array, LectinOracle again achieved a quantitative correlation to independent experimental data (Figure 3D-F), with some lectins, such as AAL and UEA-I, even showing a strong correlation between predictions and experimental observations (Pearson’s correlation coefficient > 0.7). We extended this to two additional array types, the microbe-focused glycan array^[47]^ (Figure 3G-I) and the sialic acid array^[48]^ (Figure 3J-L). In both cases, we observed moderate to strong correlations between the predicted binding values from LectinOracle and the experimentally measured binding.

In some cases, such as DC-SIGN on the microbe-focused array (Figure 3H), LectinOracle correctly predicted the binding to one cluster of glycan sequences (mannose-rich glycans) yet missed another cluster of glycans in its predictions that was measured to bind (Lewis X-type structures^[49]^). This suggests that, with more data from more diverse glycan arrays, LectinOracle can be further improved to detect all binding specificities of a lectin. Overall, this and the high precision of our predictions (Figure S7A and D) implies that any errors in LectinOracle predictions are more likely to be false negatives than false positives, raising the confidence in binding predictions. These validations across multiple independent arrays included bacterial, viral, plant, fungal, and mammalian lectins. Additionally, the majority of glycans in these custom arrays were not present in the CFG and Imperial College arrays used for training LectinOracle. We therefore concluded that LectinOracle can generalize to new lectins, new glycans, and new contexts (e.g., different linkers), as long as they are not too far removed from the training set, such as the antibodies mentioned above.

### Investigating lectomes in host-microbe interactions with LectinOracle uncovers shifting binding repertoires

Lectins are widespread throughout all kingdoms of life and carry out a panoply of essential functions, from combating pathogens^[50]^ to distinguishing self and foreign tissue^[51]^. Yet the set of lectins that have been experimentally characterized to a sufficient degree only represents a sliver of the total set of lectins in nature. Genome annotation efforts, based on sequence similarity to known lectins, have resulted in databases such as LectomeXplore^[9]^. There, researchers have catalogued predicted lectins from thousands of species, as well as their lectin class.

Seeking to better understand these predicted lectins, we used our LectinOracle platform to analyze lectins in LectomeXplore with a prediction score of 0.5 or above (i.e., at least 50% similarity to known lectins). This corresponded to 120,523 putative lectins from 7,753 species, which in turn represented 113,573 unique protein sequences. We calculated ESM-1b representations for these sequences and observed that, while many assigned lectin classes clustered together based on sequence similarity (Figure 4A), classes such as ficolin-like lectins or L-rhamnose binding lectins demonstrated a fragmented cluster behavior, suggesting a sequence diversity or sub-classes within these categories. It should be noted that broad classes such as ficolin-like lectins, defined by containing fibrinogen-like domains, could also contain non-lectin proteins.

**Figure 4.**
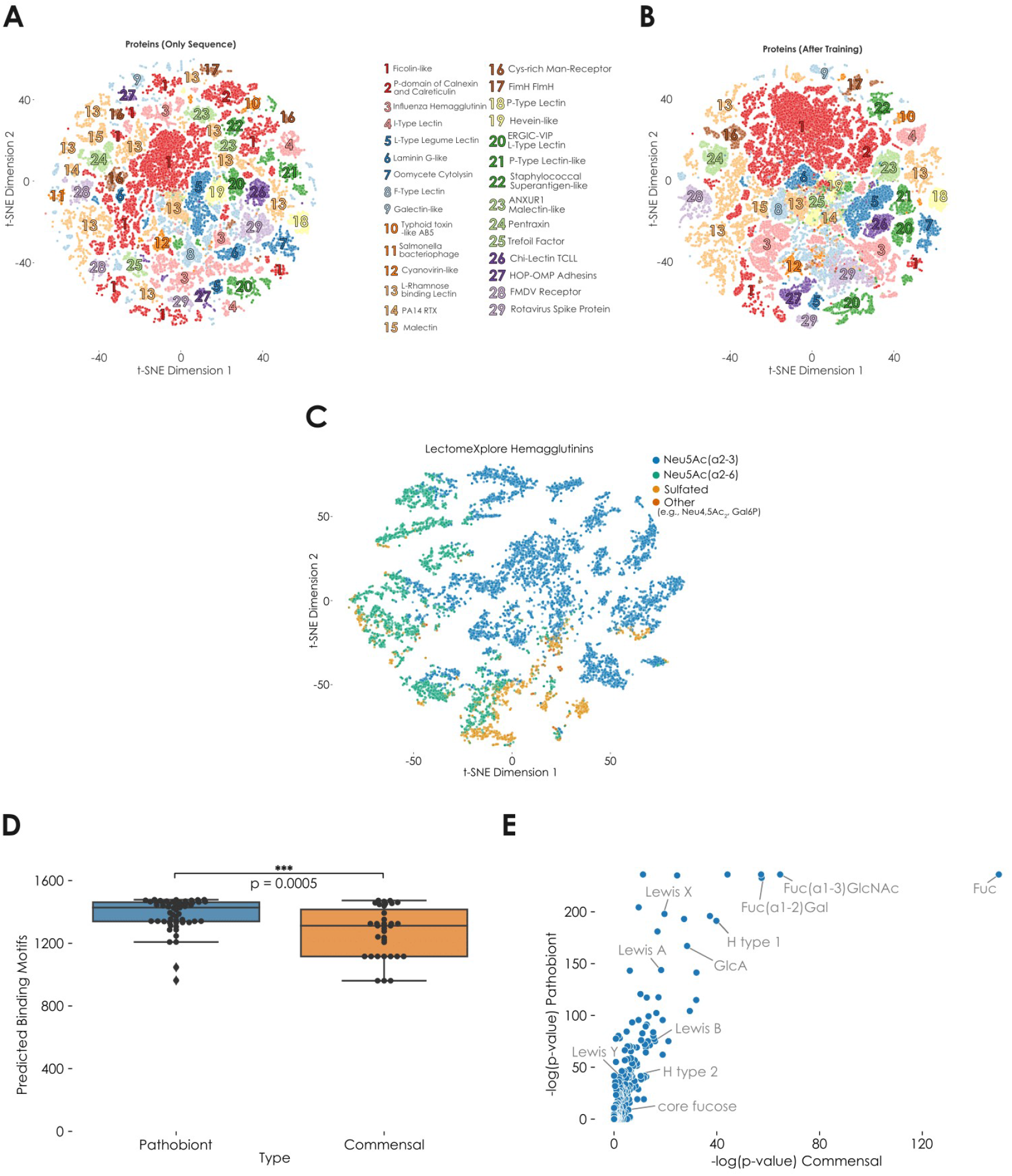
Analyzing lectomes and their role in biology with LectinOracle. **A-B)** Lectins from LectomeXplore clustered based on sequence similarity or binding specificity. Learned representations from the pre-trained ESM-1b model (**A**) or a trained LectinOracle model (**B**) were extracted for all 120,523 putative lectins in LectomeXplore with a similarity score higher than 0.5. Lectin classes with at least 400 examples are annotated in both (A) and (B). **C)** Predicted glycan-binding specificity for hemagglutinins on LectomeXplore. For 9,752 hemagglutinin sequences from LectomeXplore, we plotted their learned representation from a trained LectinOracle model and colored them based on their predicted glycan-binding specificity. Motifs were colored if they showed a significant enrichment in predicted binding, according to motif-based one-tailed Welch’s t-tests and a Holm-Šídák correction for multiple testing. **D)** Pathobionts in the vaginal microbiome have a larger predicted binding repertoire than commensals. For all strains in the dataset, we used a trained LectinOracle model to predict the binding motifs of their lectins. The number of motifs with a predicted binding above zero is shown for both pathobionts and commensals. Statistical significance between groups was established with a two-tailed Welch’s t-test. **E)** Enriched predicted binding motifs for lectins from pathobionts and commensals. For all lectins from pathobionts and commensals, we predicted their binding to a range of glycan motifs and analyzed enriched binding motifs via one-tailed Welch’s t-tests and a Holm-Šídák correction for multiple testing. The resulting p-values per motif are shown as −log(p-value), with representative motifs annotated.

Next, we visualized the learned lectin representations after training LectinOracle (Figure 4B). These representations, factoring in learned glycan-binding specificity in addition to sequence similarity, yielded an improved clustering, with a lower cluster variance for most lectin classes (Figure S8A). Classes such as Foot-and-mouth disease virus (FMDV) receptor lectins, which showed two distinct clusters when clustering by ESM-1b representations, were unified when clustering on both sequence similarity and binding specificity. We also note that the sequences in a cluster often corresponded to very uniform LectomeXplore score values (Figure S8B-C), implying that they form pools of sequences that are similar to each other, with a comparable similarity to the profile used for searching for the lectin class in genomes. Some classes, such as ERGIC VIP (Endoplasmic Reticulum Golgi Intermediate Compartment/Vesicular Integral Proteins) L-type lectins, formed distinct clusters with different score distributions, suggesting that these clusters correspond to different pools of lectins from the ERGIC VIP L-type class. This could pave the way for a more fine-grained classification of lectins that may differ in important aspects.

We then further analyzed additional lectin classes on LectomeXplore. One example of broad interest are hemagglutinin proteins, which are crucial for influenza virus cell entry, as the glycan-binding specificity of hemagglutinin can determine host range^[4]^. Classically, hemagglutinins are split between binding α2-3-linked Neu5Ac (avian host) and α2-6-linked Neu5Ac (mammalian host), with additional reports that sulfated^[52]^ and phosphorylated^[53]^ glycans may be bound. When analyzing the 9,752 hemagglutinin sequences from LectomeXplore with a score above 0.5 via LectinOracle, we could clearly separate hemagglutinins into three broad clusters, corresponding to Neu5Ac(α2-3), Neu5Ac(α2-6), and sulfated glycans as their major binding epitope (Figure 4C). We additionally noted the occurrence of smaller clusters that seemed to prefer binding to phosphorylated glycans or O-acetylated Neu5Ac (Neu4,5Ac_2_).

Further analyses into other LectomeXplore classes, such as clustering staphylococcal superantigen-like lectins or F-type lectins into sub-groups with different glycan-binding preferences supported the ability of LectinOracle to further characterize these putative lectins at scale (Figure S9). These results demonstrate that the lectin annotation pipeline could be extended to also include predicted binding specificity for further insight.

Recent work has focused on analyzing the lectomes of pathobionts and commensals in the vaginal microbiome, finding a higher number of lectins in pathobionts than in commensals^[54]^. Using information of which glycan motifs the respective lectin classes typically bind resulted in the conclusion that the lectomes from pathobionts seemed to bind to a higher diversity of glycan motifs. Using predictions from our LectinOracle model, we arrived at a similar result in that lectomes from pathobionts were predicted to bind to more glycan motifs than those from commensals (Figure 4D), which might aid pathobionts in adhering more robustly to mucosal surfaces^[55]^.

We then set out to capitalize on the predictive capabilities of LectinOracle to investigate which glycan motifs were targeted by pathobionts and commensals, respectively (Figure 4E). Based on our predictions, lectins from both pathobionts and commensals seemed to be particularly enriched in binding to fucose-containing motifs. In accordance with our earlier analysis (Figure 5A), pathobionts exhibited a greater diversity of highly enriched binding motifs, including various fucosylated motifs that are prevalent in human glycans (Lewis X, Lewis A, H type 1, etc.) and that are known to mediate adhesion of other pathobionts such as *Helicobacter pylori*^[56]^, These structures could thus also be used by the pathobionts of the vaginal microbiome to adhere to mucosal surfaces, showcasing the utility of LectinOracle to yield further insight into biological contexts involving lectins.

## Discussion

If glycosyltransferases and related enzymes could be construed as the “writers” in glycobiology, lectins would represent the “readers” of the glycocode. Since glycans dominate the surface area of most cells, protein-mediated interactions between cells or organisms typically rely on lectins that recognize specific glycan motifs. Yet very few lectins follow a strict one protein - one motif correspondence such as zinc finger proteins in DNA recognition^[57]^. Mostly, lectins span the whole range, from a narrowly defined binding specificity, such as SNA, to a broader, more relaxed binding specificity that is still well-defined, such as in the case of ConA. Knowing which glycan motifs are bound by any given lectin is a nontrivial endeavor and usually entails months or even years of dedicated study. This is a problem both for timely issues, such as ascertaining the glycan-binding specificity of a pandemic-causing pathogen^[58]^, as well as for systematic issues, as the high-throughput characterization of whole lectomes^[9]^ is currently infeasible. We view LectinOracle as a means to alleviate these bottlenecks, as well as to further characterize and cluster well-studied lectins, by obtaining binding predictions for lectin sequences in a rapid and scalable manner.

LectinOracle can be extended to both new lectins and new glycans, paving the way for its integration into the routine study and usage of lectins. Potential applications, as shown here, span the range from in-depth characterization of the binding profile of individual lectins to the analysis of whole lectomes and their binding profile in health and disease, with potential mechanistic and biomedical implications. In future work, approaches such as presented here could also be combined with the tissue-specific glycome of a host species^[59]^ to identify specific glycan receptors that could be physiologically relevant for the adhesion of a pathogen.

A potential limitation of our work could lie in extreme generalizability. We already mentioned that distinct protein subgroups, such as antibodies, might not be amenable to be used as inputs for LectinOracle if they are currently not represented in the training data. Further, while new glycans can be readily used as inputs for LectinOracle, we are currently limited to glycans composed of the monosaccharides used for training LectinOracle. Unseen monosaccharides would not have a learned representation and could, at this stage, not be interpreted by LectinOracle. Fortunately, LectinOracle was trained with a large set of 80 monosaccharides and linkages, which will enable researchers to work with most glycans. The integration of future glycan arrays with even more diverse glycans will further improve this type of generalizability.

Recent research has shed light on the differences between binding conditions on glycan arrays and physiological presentation of glycans on cells and tissues^[60–63]^, including issues such as crowding, diversity, and linker properties. For some lectins, binding preferences that were found on glycan arrays could not be recapitulated on tissues and vice versa. As LectinOracle was trained on glycan array data, these limitations could extend to its predictions as well. Yet, at this stage, there is by far not enough physiological binding data available to train any kind of model. Further, current expert inferences of binding specificity are also informed by array data and typically do not deviate extensively from physiological behavior. By extension, predictions made with LectinOracle should thus also yield physiologically relevant predictions on average. Additionally, as more structural data of lectin-glycan complexes become available, this type of information could eventually be added as a “third arm” of LectinOracle, further improving prediction results and offering a more direct path to mechanistic interpretation. However, this step should ideally be present as an optional input, to not impede generalizability.

In general, learning protein-glycan interactions from sequences instead of structures allowed us to leverage a vastly larger amount of data. Drawbacks with protein-glycan co-crystallization data, apart from data scarcity, include the lack of available quantitative binding data and the absence of negative examples. While every co-crystal represents a positive example of protein-glycan interaction, structural information alone does not provide data as to whether a lectin will conclusively not bind a certain glycan. The array data, which we used here, contains an abundance of true negative examples, of lectins not binding to certain glycans, and is exclusively composed of quantitative binding data. In contrast to treating lectins as essentially black boxes^[16]^, our sequence-based approach further allows us to extract information from amino acid sequences and extrapolate to new lectins. This middle ground approach allowed us to combine the best of both worlds and construct a highly generalizable model that enables indepth analyses of lectin binding behavior. We therefore envision that LectinOracle will be a versatile platform to advance glycobiology as well as the many other life science disciplines in which lectins exercise an important role.

## Materials & Methods

### Dataset Construction

For the lectin-glycan dataset, we manually curated data from 3,228 glycan arrays from the Consortium for Functional Glycomics database, using a custom script to extract the data from the Excel files. We also added 100 glycan arrays from the Carbohydrate Microarray Facility of Imperial College London to our dataset. For all glycan arrays, we converted glycan descriptions to IUPAC-condensed via a mapping table (Table S4). Wherever possible, we also collected meta-data about the sample, such as protein sequence, database identifiers, and expression system (Table S2).

### Data Processing

First, we averaged columns that were mapped to the same glycan sequence in IUPAC-condensed nomenclature. Then, we selected the subset of data from array experiments where the protein sequences were available to us. This resulted in a final set of 2,709 glycan array experiments for training LectinOracle. All array experiments were then normalized by Z-score transformation. Then, we averaged data from experiments using the same protein sequence in different concentrations or under different buffer conditions. We also averaged data from glycans attached to the array via different linker sequences. These procedures were carried out to enable an interaction prediction of a protein sequence to a glycan without special consideration to environmental conditions, enabling generalizability. This resulted in 564,647 unique protein-glycan interactions which we used to train and evaluate LectinOracle.

### Model Training

For model training, we split the dataset into a training (90%) and a test set (10%), ensuring that no proteins were present in both the training and the test set. Then, we converted the data in both sets into the format (protein sequence, glycan sequence, binding Z-score). For the protein sequence, we retrieved the 1,280-dimensional representation from the trained ESM-1b model. For the glycan sequence, we used the Python package glycowork^[64]^ to convert the IUPAC-condensed sequence into a graph object, as described previously^[8]^.

LectinOracle comprised a fully connected neural network that used the ESM-1b representations as input and a SweetNet-based graph convolutional neural network with node embeddings to analyze glycan graphs. Results from these two arms were concatenated and used in another fully connected part, which resulted in a multi-sample dropout scheme^[65]^ prior to the binding prediction. Immediately before the final output, a sigmoid layer was used to scale the output to range between the maximum and minimum Z-scores observed in the overall dataset. Fully connected parts of LectinOracle constituted linear layers interspersed with leaky ReLUs, dropout layers, and batch normalization layers. All linear layers were initialized via Xavier initialization^[66]^.

All models were trained with PyTorch 1.8^[67]^ and PyTorch Geometric 1.8^[68]^, using a single NVIDIA^®^ Tesla^®^ P100 GPU. We used batch sizes of 128 for both training and test sets. Final hyperparameters after optimizing via cross-validation were a starting learning rate of 0.0005, using ADAM as an optimizer, that was decayed over 80 epochs according to a cosine function. We trained LectinOracle for 100 epochs, with an early stopping criterion of 20 epochs without further improvement. For training, we used a mean squared error loss function.

### Obtaining learned glycan and protein representations

To visualize protein similarities, we either used ESM-1b representations or fine-tuned representations after training LectinOracle. For ESM-1b representations, we truncated protein sequences to a maximum of 1,000 amino acids, as ESM-1b does not support substantially longer submissions. Then, we used this as an input to the trained ESM-1b model and extracted the learned 1,280-dimensional representation from the final layer. For fine-tuned protein representations, we used this ESM-1b representation, together with an arbitrary dummy glycan sequence, as an input to LectinOracle and extracted the 128-dimensional protein representation immediately prior to concatenating protein and glycan representations (i.e., after the fully connected module that fine-tunes the ESM-1b representation to the task of predicting protein-glycan binding).

For obtaining glycan representations, we analogously used glycan sequences together with an arbitrary dummy protein ESM-1b representation as inputs to LectinOracle. Then, we extracted the 256-dimensional glycan representation immediately prior to concatenating protein and glycan representations (i.e., after the pooling operations of the graph convolutional neural network). To visualize glycan clusters for Figure 1C, we then used these glycan representations to construct a cosine distance matrix. Subsequently, we applied neighbor joining to this matrix to obtain a dendrogram based on agglomerative clustering.

### Identifying lectin binding specificity

To determine the binding specificity of a lectin, we constructed a library of disaccharides, trisaccharides, and n-1 motifs (motifs differing by one monosaccharide from full sequences in the dataset) that were observed in our glycan set. This choice of motifs was motivated by a compromise between interpretability and computational efficiency. Then, using a trained LectinOracle model, we retrieved the ESM-1b representation for the lectin and used this as input together with all motifs, to receive predicted binding scores for all motifs. Then, we subtracted the background prediction value to arrive at the final predicted binding score. For the background prediction correction, we calculated predicted binding scores for all lectins with all glycan motifs. Then, we calculated the median prediction for each motif across all lectins and considered this the prediction background (Table S5), with the assumption that no motif should be bound by all or the majority of lectins in our dataset.

## Data Availability

All code used in this study can be found at https://github.com/BojarLab/LectinOracle and all data that we used is available in the supplementary tables.

## Author Contributions

D.B. conceived the method. D.B. and J.L. performed the experiments. D.B., J.L., and E.K. prepared the dataset. All authors wrote and edited the manuscript.

