## Supplemental Figures for "LectinOracle – A Generalizable Deep Learning Model for Lectin-Glycan Binding Prediction"

### Supplementary Figures

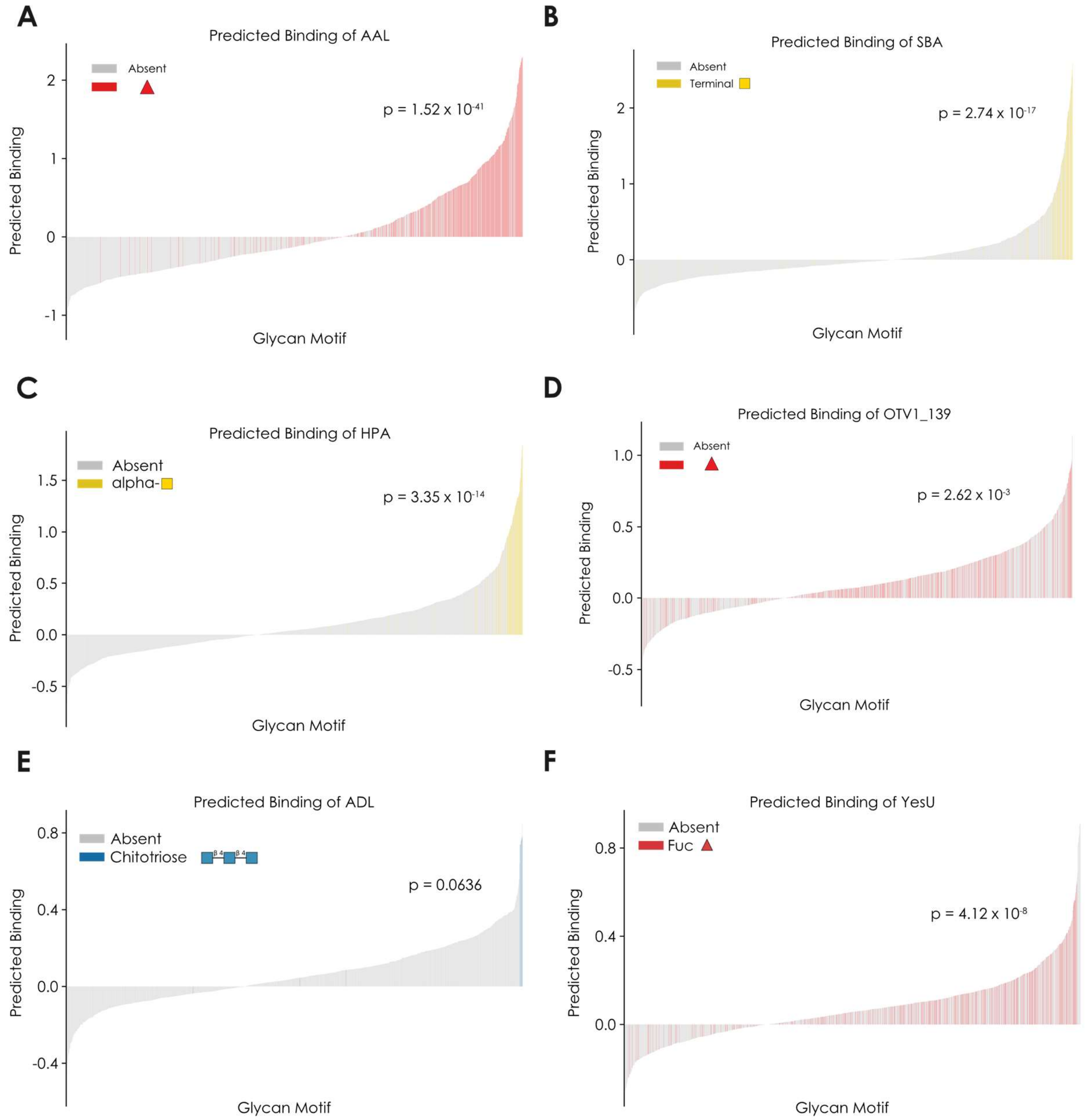


**Figure S1. Glycan-binding specificity of various lectins with LectinOracle. A-F)** For a range of glycan motifs, a trained LectinOracle model was used to predict their binding to the lectins AAL (A), SBA (B), HPA (C), OTV1_139 (D), ADL (E), and YesU (F) with their literature-annotated binding motifs or enriched binding motifs colored in.

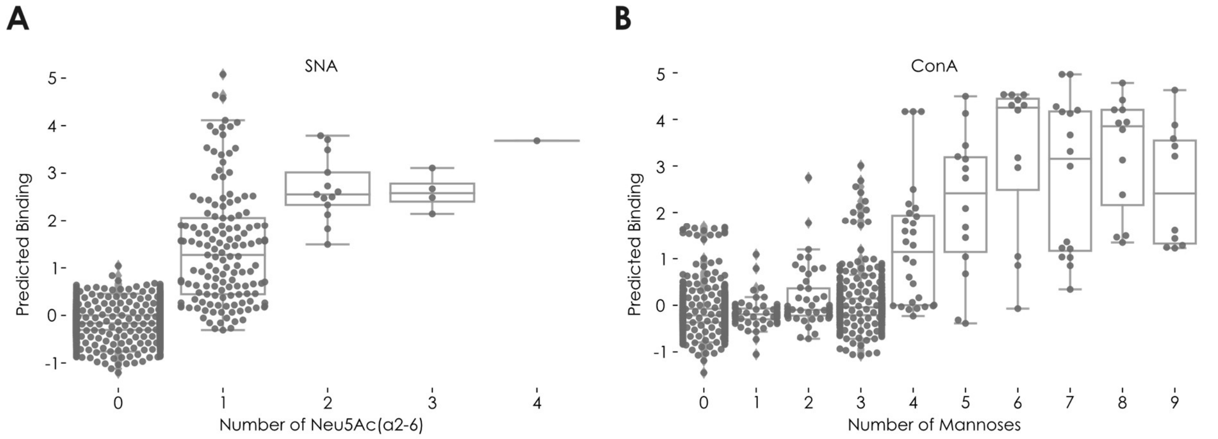


**Figure S2. Correlation of binding predictions with the number of binding epitopes. A-B)** For the lectins SNA (A) and ConA (B), we used a trained LectinOracle model to obtain binding predictions for a range of glycan motifs and full glycans. Then, we counted the occurrence of the literature-defined binding motif for the respective lectin in each glycan motif and plotted the predicted binding as a box plot.

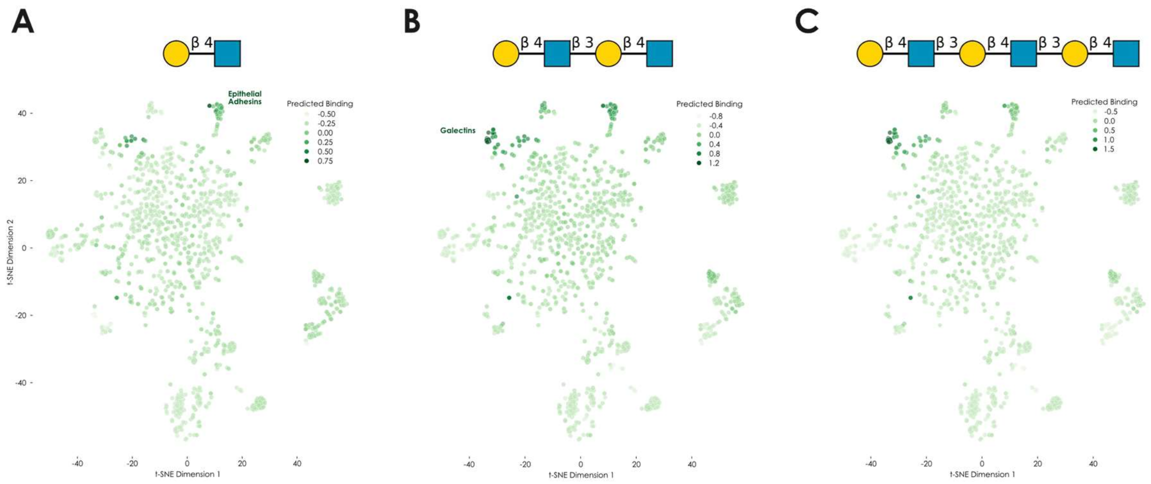


**Figure S3. LectinOracle can distinguish repeats of a disaccharide motif. A-C)** For the motifs type II LacNAc, di-LacNAc, and tri-LacNAc, we obtained the predicted binding from a trained LectinOracle model for all lectins and colored the learned representation from Figure 2B accordingly. Clusters of lectins exhibiting elevated predicted binding to at least one motif are annotated with enriched lectins.

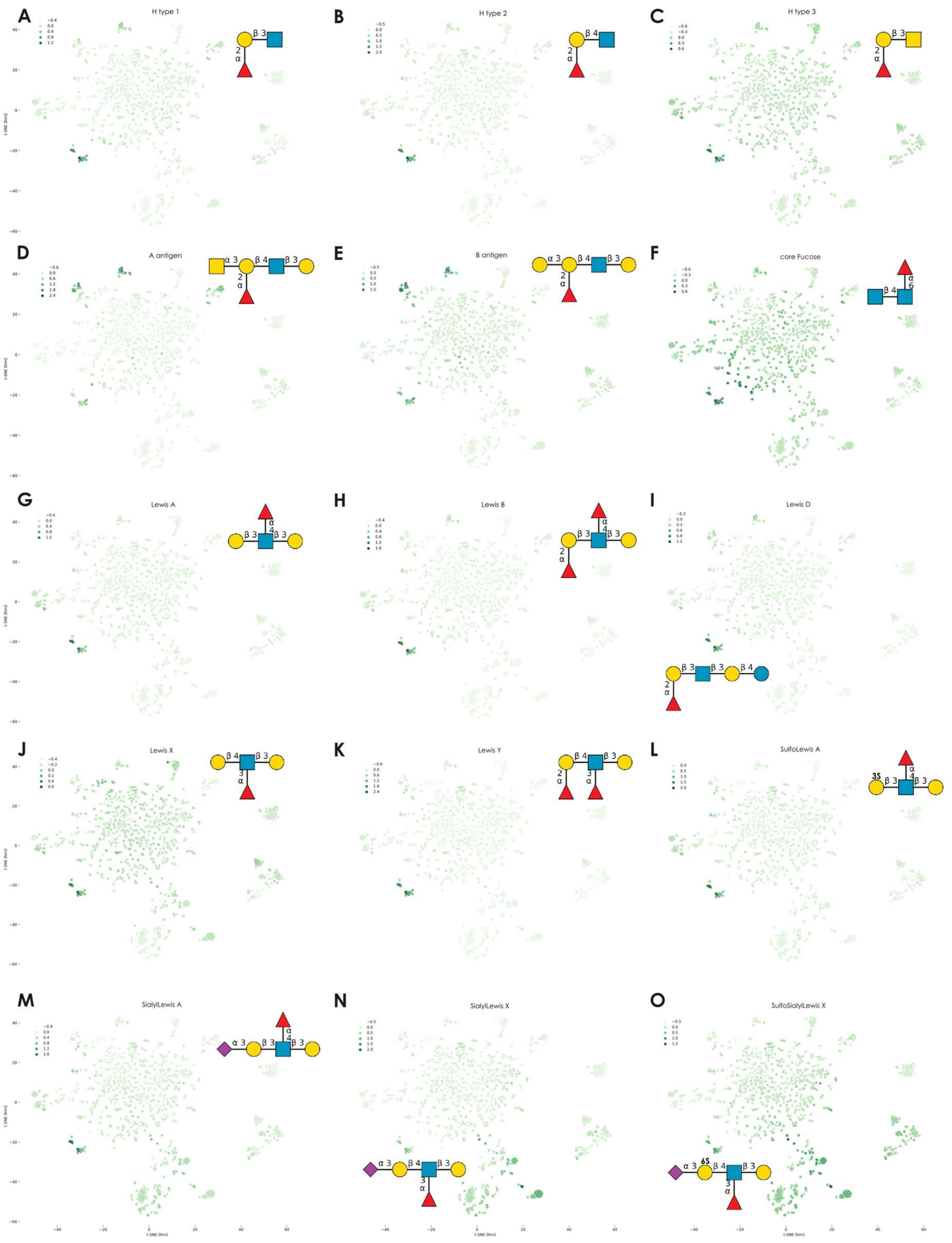


**Figure S4. Predicted binding of lectins to important glycan motifs. A-O)** We used a trained LectinOracle model to infer the binding of all lectins in our dataset to a range of important glycan motifs (A-O) and then colored the t-SNE visualized lectin representation (see Figure 2B) according to the predicted binding to the respective glycan motif. The glycan motifs are shown for each panel in SNFG format.

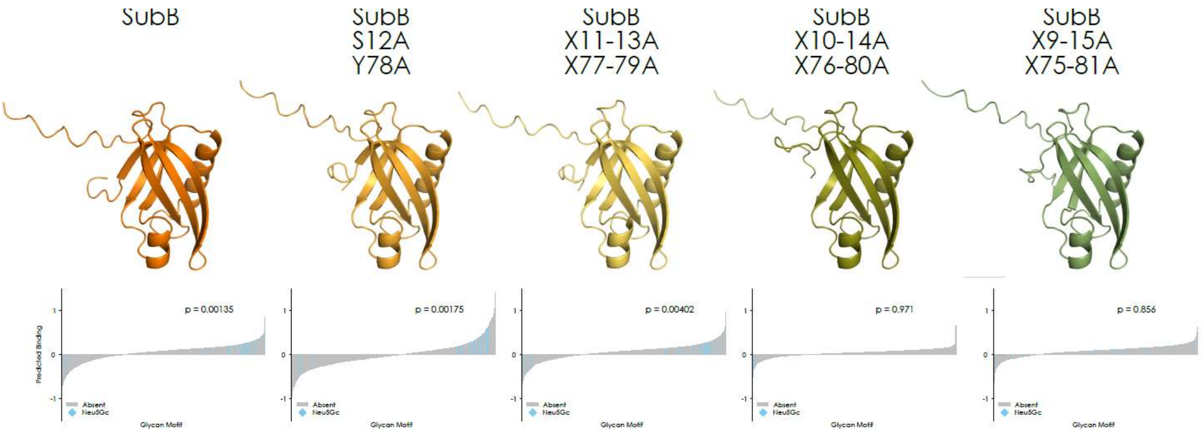


**Figure S5. Sensitivity of LectinOracle predictions for SubB - Neu5Gc interactions.** Similar to Figure 2C, we retrieved the binding predictions for a range of glycan motifs for SubB and various alanine-substitution mutants from a trained LectinOracle model. Binding predictions are colored by the presence or absence of Neu5Gc and the enrichment p-values were calculated via a one-sided Wilcoxon signed-rank test. We note that there are far fewer Neu5Gc-containing motifs than for Neu5Ac, due to a dearth of Neu5Gc-containing glycans on the glycan array we used to train LectinOracle. Protein structure predictions made with AlphaFold2 are shown for the wild-type protein and each mutant.

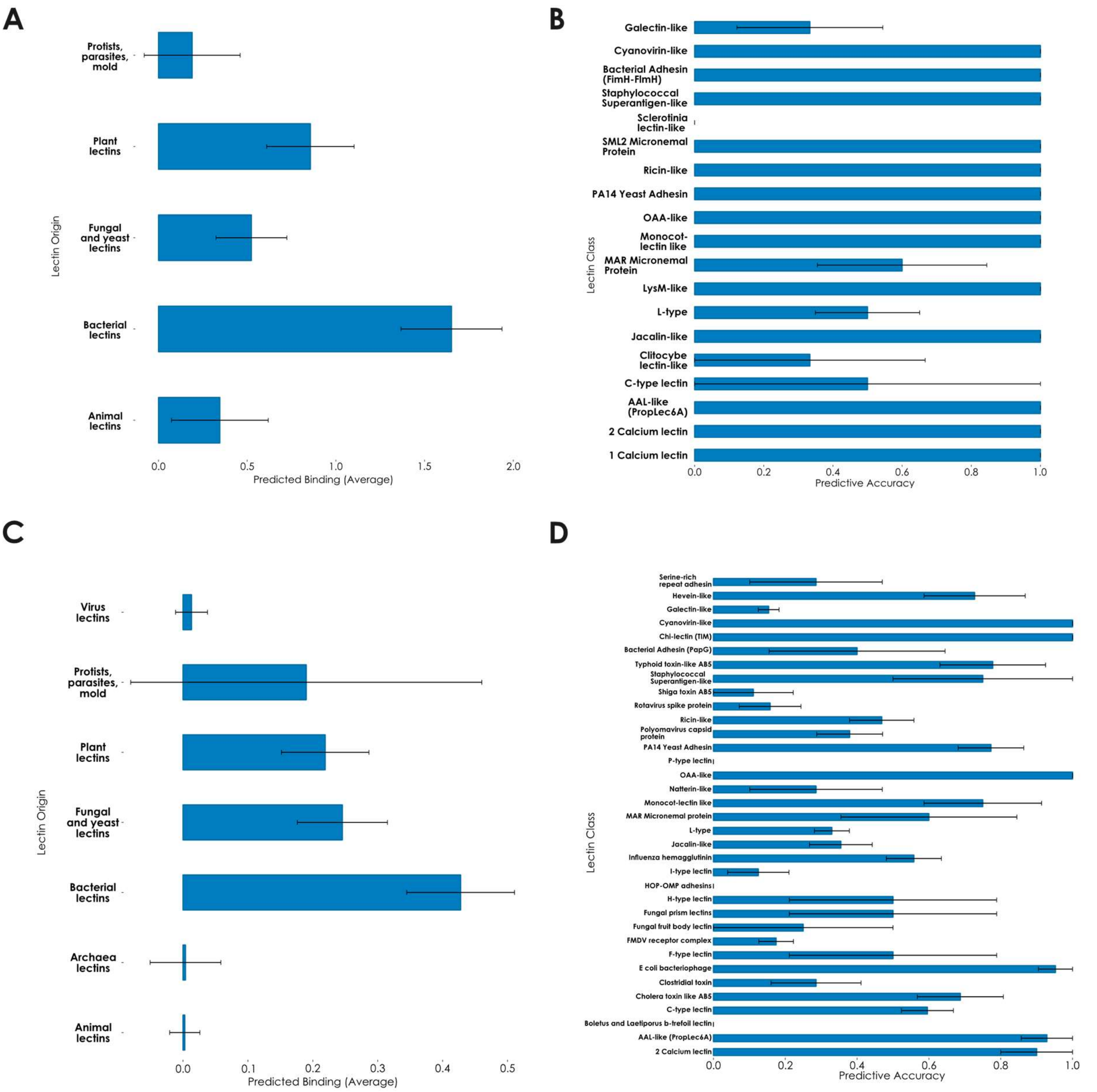


**Figure S6. Comparing LectinOracle predictions and UniLectin3D crystal structures. A-B)** Predictions for UniLectin3D proteins contained in our dataset. For all lectin-glycan structures for which the lectin was part of our dataset, we used a trained LectinOracle model to predict the binding of the lectin-glycan pair and show the average predicted binding grouped by lectin origin (A) or class (B). **C-D)** Predictions for all UniLectin3D proteins. For all lectin-glycan structures, we used a trained LectinOracle model to predict the binding of the lectin-glycan pair and show the average predicted binding grouped by lectin origin (C) or class (D), for classes with at least three proteins in UniLectin3D. Shown are means ± s.e.m.

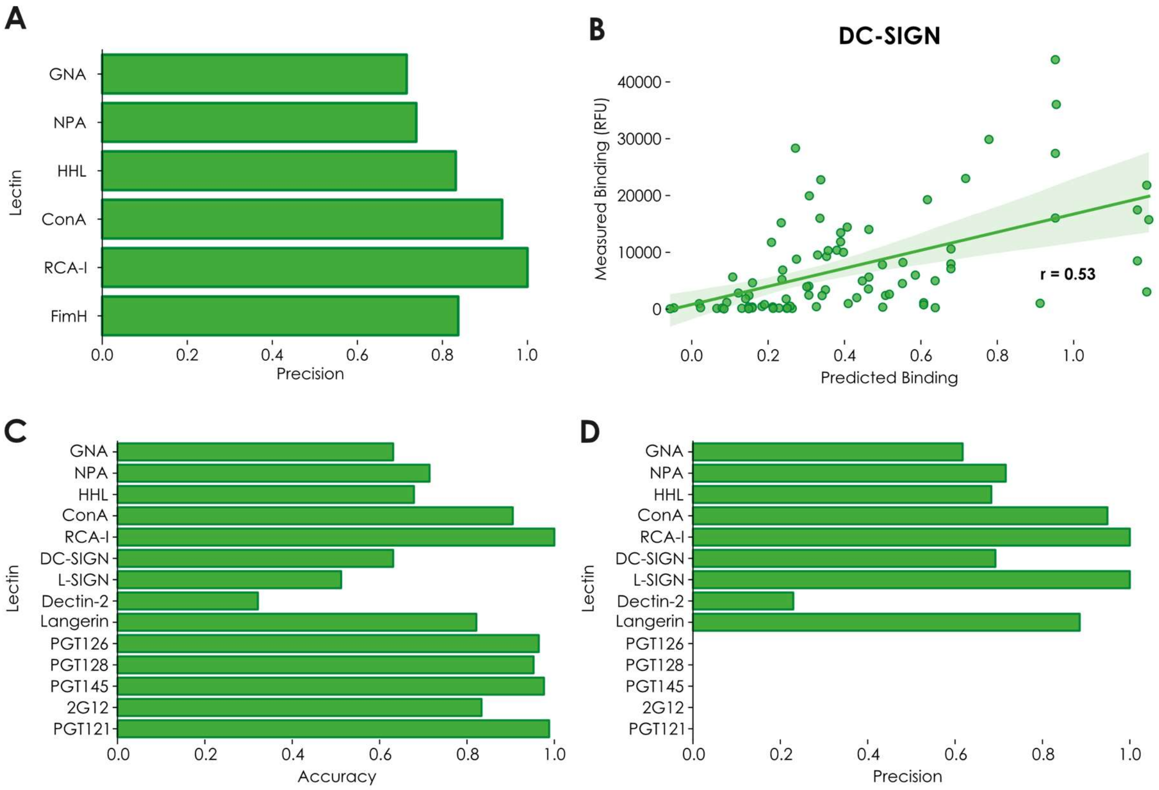


**Figure S7. Additional validation of LectinOracle with data from the oligomannose array. A)** Precision of LectinOracle predictions for lectins tested on the oligomannose array. For each lectin-glycan pair, we assigned it the label "bound" or "predicted bound" if the observed relative fluorescence units (RFU) were at least 10% of the maximum RFU or if the predicted binding was at least 0.5, respectively. **B)** Correlating experimentally observed binding with predictions for the lectin DC-SIGN. Correlations between experimental data and predictions were done via fitting a linear regression and r represents Pearson's correlation coefficient. **C-D)** Validating LectinOracle on another set of lectins tested on the oligomannose array. For a set of plant lectins, mammalian lectins, and antibodies, we compared LectinOracle predictions similar to (A) and depict predictive accuracy (C) and precision (D).

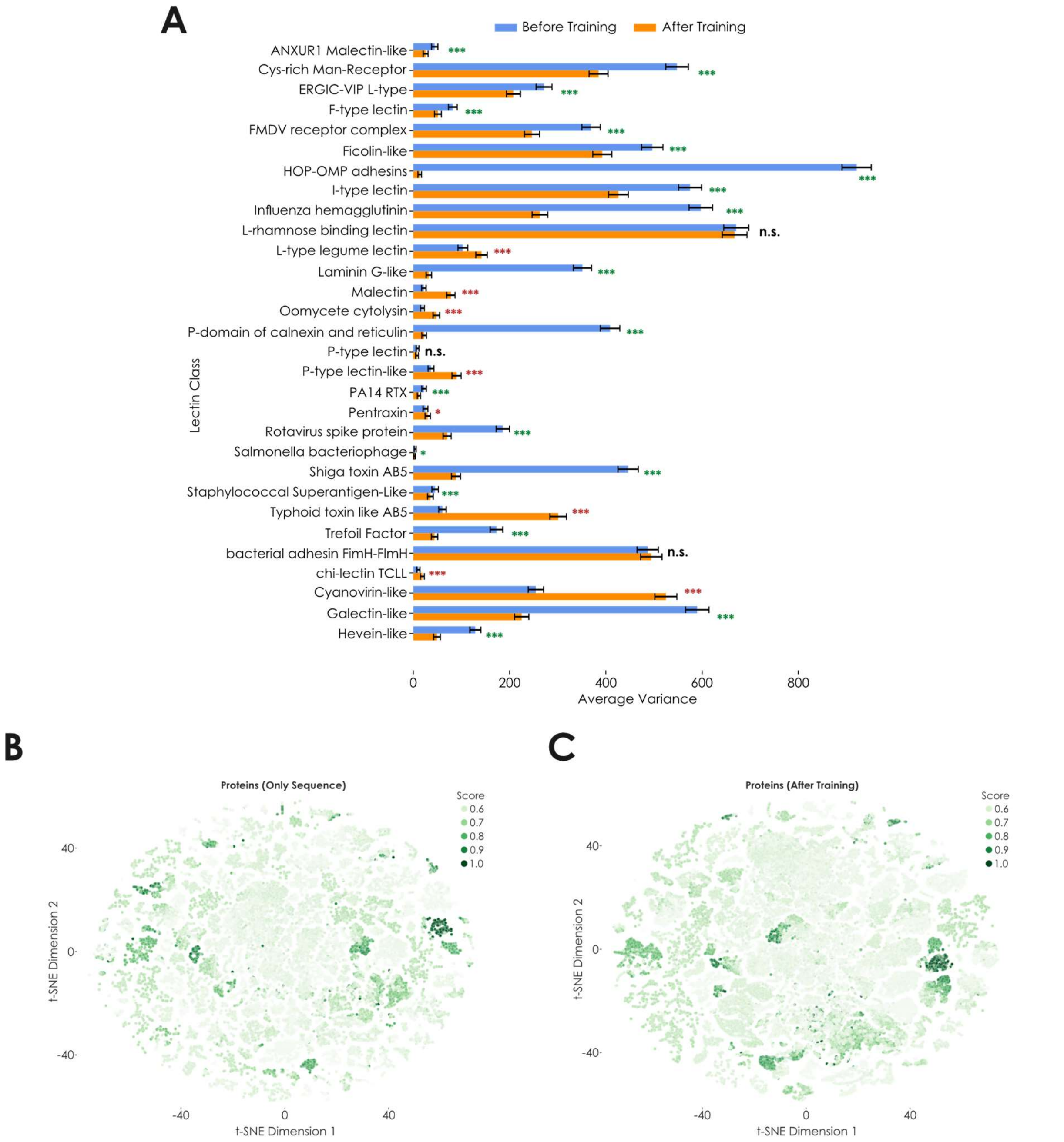


**Figure S8. Analyzing putative lectins in LectomeXplore. A)** Training LectinOracle improves clustering by lectin class. Variance by lectin class is shown before and after training, with significant differences being tested via an F-test. * $p < 0.05$, ** $p < 0.01$, *** $p < 0.001$. **B-C)** Lectins from LectomeXplore clustered based on sequence similarity or binding specificity. Learned representations from the pre-trained ESM-1b model (B) or a trained LectinOracle model (C) were extracted for all 120,523 putative lectins in LectomeXplore with a similarity score higher than 0.5. Lectins were colored by the sequence similarity score from LectomeXplore.

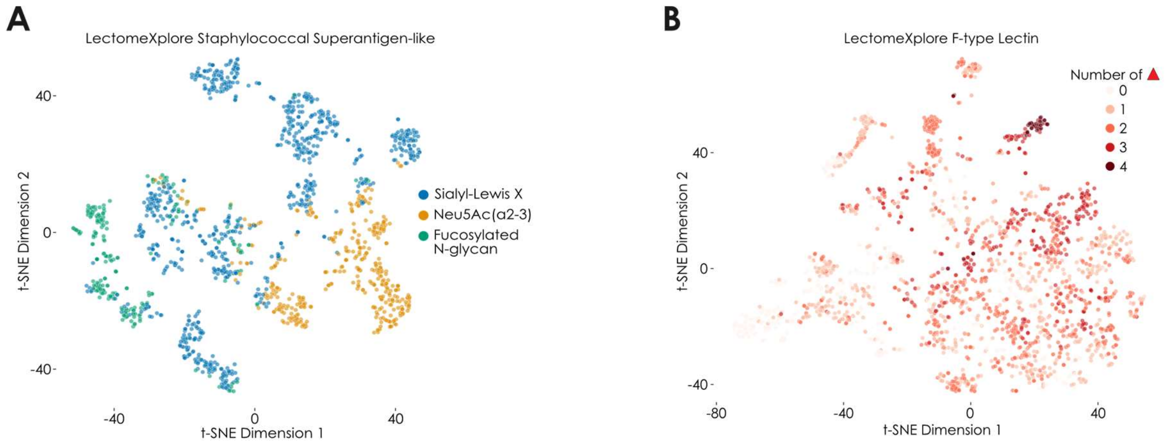


**Figure S9. Analyzing LectomeXplore lectin classes with LectinOracle. A-B)** We used LectinOracle to predict the binding specificity for all 1,411 staphylococcal superantigen-like (SSL, A) or 2,206 F-type (B) lectins with a score above 0.5 in LectomeXplore. Shown are the learned lectin representations learned by LectinOracle, analogous to Figure 4C, colored in by preferred binding motif. Lectins colored in for "Neu5Ac(α2-3)" binding did not show preferred binding to Sialyl-LewisX motifs. For (B), we counted the number of fucoses in the top five enriched motifs as a measure of fucose binding promiscuity
